# Mitochondrial Ca^2+^ controls pancreatic cancer growth and metastasis by regulating epithelial cell plasticity

**DOI:** 10.1101/2024.08.08.607195

**Authors:** Jillian S. Weissenrieder, Jessica Peura, Usha Paudel, Nikita Bhalerao, Natalie Weinmann, Calvin Johnson, Maximilian Wengyn, Rebecca Drager, Emma Elizabeth Furth, Karl Simin, Marcus Ruscetti, Ben Z. Stanger, Anil K. Rustgi, Jason R. Pitarresi, J. Kevin Foskett

## Abstract

Endoplasmic reticulum to mitochondria Ca^2+^ transfer is important for cancer cell survival, but the role of mitochondrial Ca^2+^ uptake through the mitochondrial Ca^2+^ uniporter (MCU) in pancreatic adenocarcinoma (PDAC) is poorly understood. Here, we show that increased MCU expression is associated with malignancy and poorer outcomes in PDAC patients. In isogenic murine PDAC models, *Mcu* deletion (*Mcu*^KO^) ablated mitochondrial Ca^2+^ uptake, which reduced proliferation and inhibited self-renewal. Orthotopic implantation of MCU-null tumor cells reduced primary tumor growth and metastasis. *Mcu* deletion reduced the cellular plasticity of tumor cells by inhibiting epithelial-to-mesenchymal transition (EMT), which contributes to metastatic competency in PDAC. Mechanistically, the loss of mitochondrial Ca^2+^ uptake reduced expression of the key EMT transcription factor Snail and secretion of the EMT-inducing ligand TGFβ. Snail re-expression and TGFβ treatment rescued deficits in *Mcu*^KO^ cells and restored their metastatic ability. Thus, MCU may present a therapeutic target in PDAC to limit cancer-cell-induced EMT and metastasis.

## Introduction

Pancreatic cancer is one of the most lethal cancers in the United States, with the most common form, Pancreatic Ductal Adenocarcinoma (PDAC), having a five-year survival rate of only ∼13%^1–3^. Patient treatment is hampered by late diagnosis, early metastasis, and poor treatment responses. The vast majority of PDAC patients present with metastatic disease, which is often the cause of death^4^. PDAC is generally heterogeneous, refractory to most treatments, and driven by currently un-targetable driver mutations, though some progress has been made with specific *KRAS* mutations^5^. Thus, while targeted therapies have vastly improved survival in many other malignancies, such as breast and prostate cancer, the standard of treatment for PDAC remains resection and cytotoxic chemotherapy^1,5^.

Genetic regulators of the metastatic cascade remain largely undefined, suggesting that non-genetic cellular plasticity contributes to the underlying biological processes driving metastasis. Such plasticity changes tumor cell biology to alter metabolic requirements, responses to chemotherapeutics, growth rates, and responses to the immune system to promote cell survival, growth, and metastasis. Given the early dissemination of tumor cells in PDAC development, cellular plasticity is of particular interest. Epithelial-to-mesenchymal transition (EMT) has emerged as a means for tumor cells to gain pro-metastatic features through a progressive loss of epithelial markers, such as E-cadherin, and an increase in mesenchymal markers, like N-cadherin ^6–10^. Classically, this transition is mediated by multiple transcription factors, including Snail, Slug, and Twist^6,7,9,10^ downstream of signaling pathways including TGFβ^8,11,12^. These EMT transcription factors actively repress the epithelial program and promote a pro-invasive mesenchymal phenotype that facilitates metastasis. Recently, lineage-labeled genetically engineered mouse models (GEMMs) of PDAC have revealed that partial or hybrid EMT states, where tumor cells co-express both epithelial and mesenchymal genes, are prevalent in pancreatic cancer ^13^. Distinct from classical or complete EMT, these partial EMT states are regulated by protein localization, metabolism, and second messenger signaling ^13–17^. Notably, EMT has been linked to metabolic alterations and enhanced Ca^2+^ signaling ^6,12,18–21^. Despite observations that cytoplasmic Ca^2+^ promotes EMT, little is known regarding the mechanisms that link intracellular Ca^2+^ homeostasis to EMT.

While Ca^2+^ signaling is known to strongly affect many cancer-related phenotypes, the role of mitochondrial Ca^2+^ signaling in PDAC is poorly understood^12^. Ca^2+^ signaling in other cancers has been implicated in pro-tumor phenotypes, including therapeutic resistance and modulation of cellular identity through plasticity events such as EMT ^12,22–24^. Many cancer cells appear to be “addicted” to Ca^2+^ flux from the endoplasmic reticulum (ER) to mitochondria, which may represent a therapeutic vulnerability ^25^. Canonically, this signaling occurs at mitochondria-associated membranes (MAMs), where Ca^2+^ released by ER-localized inositol 1,4,5-trisphosphate receptors (IP3Rs) is taken up by the mitochondrial Ca^2+^ uniporter channel complex (MCU). Critically, when MCU is lost, this uptake does not occur ^26^. The Ca^2+^ released by IP3Rs is rapidly taken up by MCU in a quasi-synaptic manner at MAMs due to the high electrochemical gradient of the mitochondrial inner membrane (membrane potential ∼ -150 to -180 mV) and the close apposition of ER and mitochondrial membranes at these sites (10-25 nm)^2,27^. This close apposition allows for ER Ca^2+^ release by constitutive low-level openings of IP3Rs and their activation in response activation of phospholipase C-coupled receptors to result in changes of mitochondrial [Ca^2+^] that regulate mitochondrial function. Previous reports have suggested that PDAC cells may depend on this flux to resist metabolic stress, since loss of MCU creates a dependency on cystine in human PDAC cells through an antioxidant-related pathway^25,28^. Here, we provide evidence that targeting mitochondrial Ca^2+^ uptake has therapeutic value in PDAC. We observe profound effects of MCU expression on PDAC tumor cell plasticity, survival, growth, and metastasis *in vivo* and *in vitro,* and elucidate a novel relationship between MCU and EMT.

## Results

### MCU is upregulated in human and murine pancreatic cancers

We examined tissue and publicly available data sets to identify links between MCU expression, tumorigenesis, and patient outcomes. Consistent with previously reported oncogenic functions for MCU in other cancers^25,26,28–33^, MCU protein expression is highly upregulated in PDAC tumor cells compared with normal tissue (Fig. 1A), and higher *MCU* gene expression is associated with poorer survival outcomes in the TCGA-PAAD (The Cancer Genome Atlas – Pancreatic Adenocarcinoma) cohort (Fig. 1B). Higher *MCU* expression in pancreatic tissue is correlated with *KRAS* mutations, the most common driver mutations in PDAC (Fig. 1C). Human PDAC cell lines show faster rates of mitochondrial Ca^2+^ uptake compared with normal Human Pancreatic Ductal Epithelial (HPDE) control cells (Fig. 1D-E), indicating increased MCU activity in PDAC. These findings are consistent with previous reports suggesting that cancer cells may be addicted to ER-to-mitochondrial Ca^2+^ uptake ^25^ and that they may be more tolerant of higher mitochondrial [Ca^2+^], with implications for apoptosis resistance^12^. Together, these support the notion that MCU is a putative oncogenic driver that may facilitate tumorigenesis in PDAC patients.

**Fig. 1.**
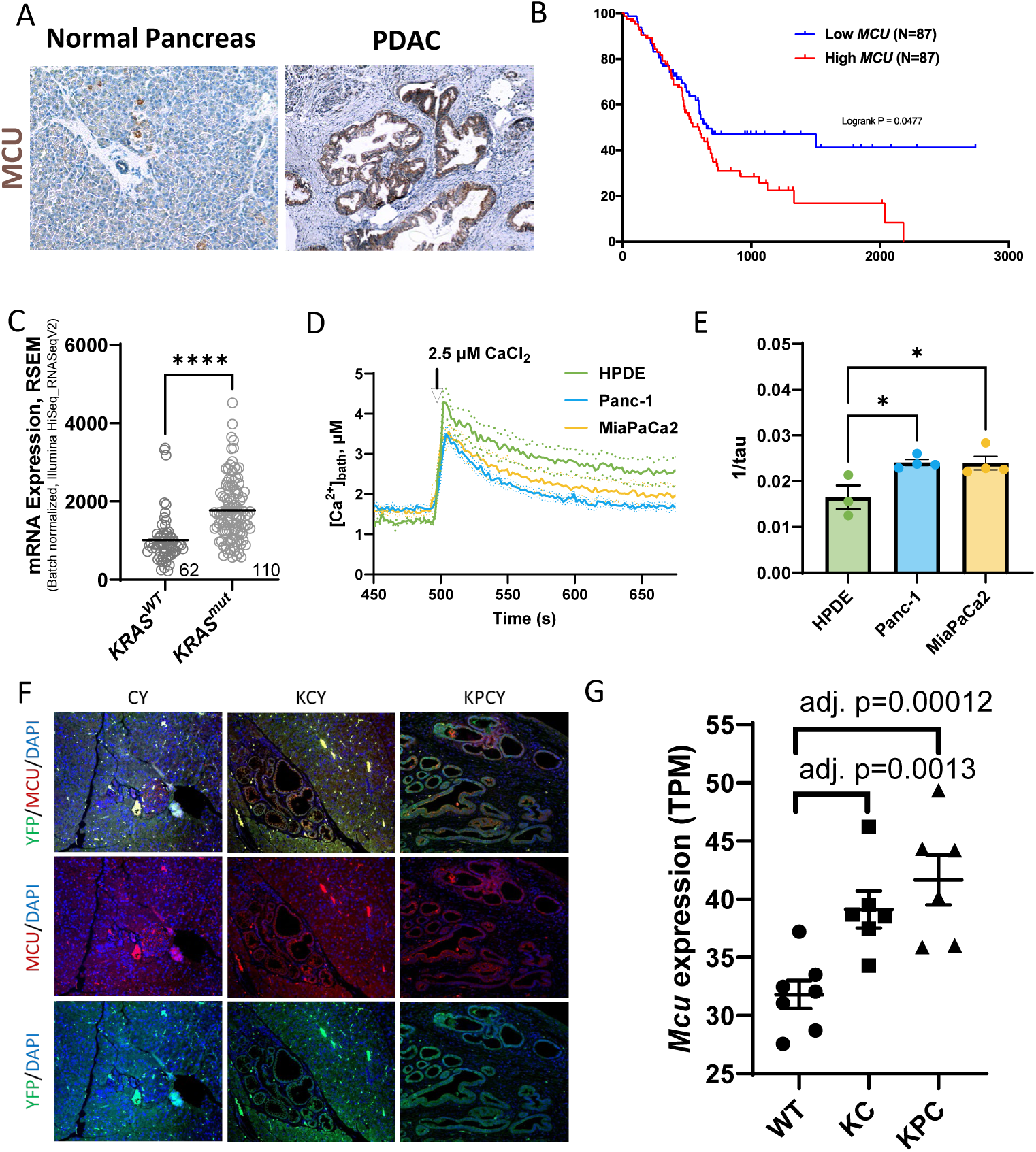
MCU expression and function are associated with malignancy in human and murine PDAC. A, MCU is highly expressed in human PDAC tissue but not in normal pancreas, as seen by immunohistochemistry. B, high *MCU* mRNA expression is associated with poor survival outcomes in the TCGA-PAAD cohort. Cohorts were split at the 50^th^ percentile, and logrank p=0.0477 via Kaplan Meier survival analysis. C, high *MCU* mRNA expression is associated with Kras mutations in the TCGA-PAAD cohort, obtained from cBioPortal. D, mitochondrial Ca^2+^ uptake is more rapid and complete in human cancer cell lines Panc-1 and MiaPaCa-2, compared with “normal” HPDE control cells. Assays carried out in biological triplicate. E, quantification of mitochondrial Ca^2+^ uptake rates in D. F, immunofluorescence imaging of tissues from normal-type CY mice, PanIN lesion-developing KCY mice, and PDAC tumor-bearing KPCY mice show high MCU expression in KCY and KPCY tissues, particularly in tumor lesions of KPCY. G, *Mcu* mRNA expression is increased in KC and KPC organoids over wild-type ductal organoids in a publicly available dataset from Tuveson et al. (GSE: 63348). Statistical analysis of survival is by Kaplan Meier analysis, while 2 group analyses were carried out with Student’s t-test. Data with 3 groups were analyzed with one-way ANOVA with Dunnett’s posthoc. *, p≤0.05. **, p≤0.01. ***, p≤0.001.

To gain a deeper understanding of when *Mcu* expression is turned on during PDAC progression, we stained for MCU in a mutant Kras- and gain-of-function Tp53-driven PDAC genetically-engineered mouse model (***<u>K</u>****ras^LSL-G12D/+^; Tr**<u>p</u>**53 ^LSL-R172H/+^; Pdx1-**<u>C</u>**re; R26^LSL-Yfp/LSL-**Y**fp^*, ‘KPCY’ mice). This mouse model is driven by mutations in *Kras* and *Trp53*, the two most common driver mutations in human PDAC that are mutated in 90% and 75% of patients, respectively^1^. The Cre-inducible Rosa26-LSL-*Yfp* allele labels tumor cells of the *Pdx1* lineage, enabling identification of PDAC tumor cells of epithelial origin^16^. Normal (exocrine acinar and endocrine islet) cells derived from *Pdx1**<u>C</u>**re; R26^LSL-**Y**fp^* (or ‘CY’) mice express appreciable levels of MCU (Fig. 1F), consistent with previous reports ^26,34,35^. MCU expression is upregulated in pancreatic intra-epithelial neoplasia (PanIN) lesions from ***<u>K</u>****ras^G12D/+^; Pdx1-**<u>C</u>**re; R26^LSL-**Y**fp/LSL-Yfp^* (KCY) mice and in YFP^+^ PDAC tumor cells from KPCY mice (Fig. 1F). Notably, YFP-negative stromal cells from KPCY mice express less MCU (Fig. 1F) than YFP-positive tumor cells (Fig. 1F), demonstrating that tumor cells upregulate MCU during tumorigenesis. Consistently, *Mcu* mRNA expression was significantly elevated with increased malignancy in a previously published RNAseq dataset [GSE63348]^36^ comparing organoids developed from the pancreas of WT, KC, and KPC mice (Fig. 1G). Taken together, these data demonstrate that MCU expression and mitochondrial Ca^2+^ uptake are upregulated in tumor cells from human PDAC patients, a phenomenon that is recapitulated in the KPCY murine model of PDAC.

### MCU promotes malignant properties of PDAC cells *in vitro*

Since MCU expression is associated with malignant phenotypes in human and murine models, we employed isogenic murine models of *Mcu^KO^* to assess the role of mitochondrial Ca^2+^ signaling in pancreatic cancer development, growth, and metastasis. Knockout of this single gene prevents the function of the MCU complex, ablating Ca^2+^ uptake into the mitochondria in response to increases in Ca^2+^ near the channel^26^. Cell lines were generated from ***<u>K</u>****ras^G12D/+^; T**<u>p</u>**53^R172H/+^; Pdx1**<u>C</u>**re; R26^LSL-**Y**fp/LSL-Yfp^; Mcu^loxP/loxP^* (KPCY-*Mcu*^Cre-KO^) mice and MCU was re-expressed at physiologically relevant levels (i.e. KPCY-*Mcu*^rescue^; Fig. 2A). The fidelity of this knockout and re-expression system was verified by Western blot analysis of clonal cell lines from each genotype (Fig. 2B). KPCY-*Mcu*^rescue^ cells express V5- and His-tagged MCU at similar levels to endogenous MCU from a previously generated KPCY murine tumor cell line, 2838.c3^37^. As expected, mitochondria in KPCY-*Mcu*^Cre-KO^ failed to take up Ca^2+^, in contrast to those in KPCY-*Mcu*^rescue^ cells (Fig. 2C), indicating that re-expressed MCU is functional. KPCY-*Mcu*^Cre-KO^ cells had reduced proliferation rates compared with KPCY-*Mcu*^rescue^ cells (Fig. 2D, ∼50% reduction), as well as reduced wound healing (Fig. 2E), spheroid formation (Fig. 2F), transwell migration (Fig. 2G) and transwell invasion (Fig. 2H). Strikingly, KPCY-*Mcu*^Cre-KO^ cells were nearly incapable of forming spheroids in anchorage-independent growth conditions, suggesting a lack of self-renewal capacity.

**Fig. 2.**
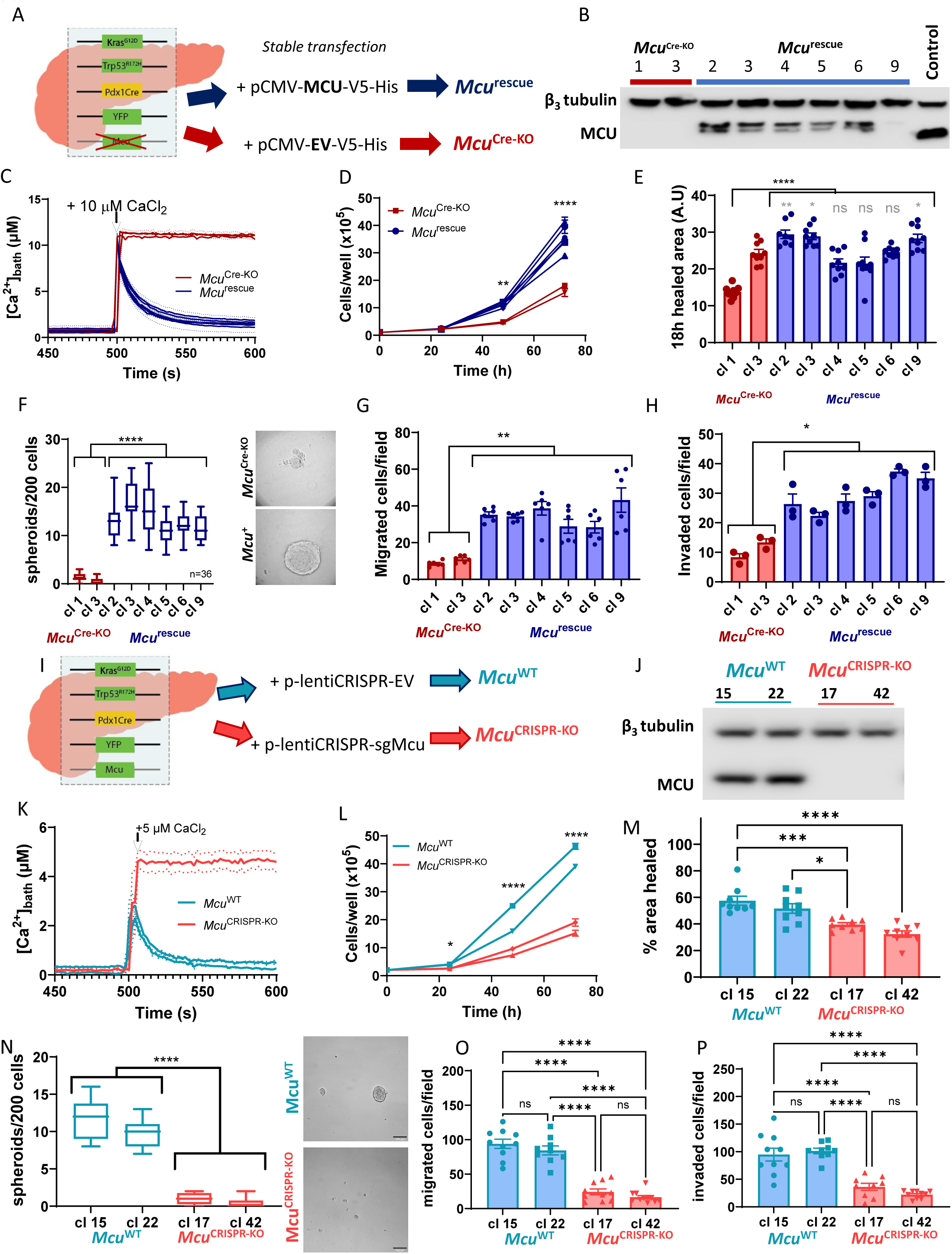
Genetic deletion of *Mcu* inhibits mitochondrial Ca^2+^ uptake and reduces growth and motility phenotypes *in vitro.* A, schematic of the development of isogenic *Mcu*^Cre-KO^ and *Mcu*^rescue^ cell lines from KPCY-*Mcu*^Cre-KO^ murine ductal cells. B, *Mcu*^Cre-KO^ clones express no MCU that is restored by stable re-expression of MCU-V5-His; 2838.c3, a KPCY-*Mcu*^WT^ cell line, is included as positive control. Pair of bands seen in *Mcu*^rescue^ cell lines due to partial degradation of the His tag. C, mitochondrial Ca^2+^ uptake is ablated in *Mcu*^Cre-KO^ cells and restored by stable expression of *Mcu*. D, *Mcu*^Cre-KO^ cells proliferate more slowly than paired *Mcu*^rescue^ isogenic cell lines. E, 18-h wound-healing assay of *Mcu*^Cre-KO^ and *Mcu*^rescue^ cells (20x). F, anchorage-independent spheroid formation. *Mcu*^Cre-KO^ spheroids, when present, are small and misshapen, in contrast to large, smooth-textured *Mcu*^rescue^ spheroids. G, migration activity in a 24-h transwell assay with FBS-containing media as chemoattractant. H, ECM-invasion capacity in 24-h transwell assay using FBS as a chemoattractant. I, schematic of the development of isogenic *Mcu*^CRISPR-KO^ and *Mcu*^WT^ cell lines from KPCY murine ductal cells. J, *Mcu*^CRISPR-KO^ clones express no MCU, in contrast to *Mcu*^WT^ isogenic controls. K, mitochondrial Ca^2+^ uptake in *Mcu*^WT^ cells is ablated in *Mcu*^CRISPR-KO^ cells. L, *Mcu*^CRISPR-KO^ cells proliferate more slowly than *Mcu*^WT^ isogenic cell lines. M, 24-h wound-healing assay of *Mcu*^CRISPR-KO^ and *Mcu*^WT^ cells. N, anchorage-independent spheroid formation. *Mcu*^Cre-KO^ spheroids, when present, are small, fragmented, and irregular, compared with large, smooth-textured *Mcu*^WT^ spheroids. O, transwell migration activity. P, ECM invasion activity. N>3 per experiment. Cell count experiments analyzed with two-way ANOVA with Tukey’s posthoc. One-way ANOVA with Sidak’s posthoc was employed for all other experiments. *, p≤0.05. **, p≤0.01. ***, p≤0.001. ****, p≤0.0001.

To ensure that these observed phenotypes were not due to compensatory mechanisms in response to the *in vivo* knockout of MCU, we also employed an isogenic CRISPR-KO model of a KPCY-*Mcu*^WT^ cell line, 2838.c5 (Fig. 2I). KPCY-*Mcu*^CRISPR-KO^ cells expressed no MCU protein (Fig. 2J) and lacked mitochondrial Ca^2+^ uptake (Fig. 2K). Consistent with the Cre-mediated *Mcu* knockout and rescue models, the deletion of *Mcu* with CRISPR strikingly reduced proliferation (Fig. 2L), wound healing (Fig. 2M), spheroid formation (Fig. 2N), migration (Fig. 2O) and invasion (Fig. 2P). Therefore, we conclude that inhibition of mitochondrial Ca^2+^ uptake by MCU knockout strongly reduces *in vitro* phenotypes associated with malignancy, metastasis, and invasion in PDAC.

### MCU promotes tumor growth and metastasis in murine xenografts

To further examine the function of mitochondrial Ca^2+^ signaling in PDAC development, the behaviors of these isogenic cell lines were interrogated *in vivo* in orthotopic implantation models of PDAC. Despite being proliferative *in vitro*, KPCY-*Mcu*^Cre-KO^ cells failed to form primary tumors after orthotopic implantation into the pancreas of C57BL/6 mice, in striking contrast to KPCY-*Mcu*^rescue^ cells (Fig. 3A-C). Notably, whereas YFP^+^ liver metastases were observed in 80% of animals implanted with KPCY-*Mcu*^rescue^ cell lines, none were observed in mice implanted with KPCY-*Mcu*^Cre-KO^ cells (Fig. 3A and Fig. 3D-F). No differences were observed in overall body weight between the two cohorts (S2A). KPCY-*Mcu*^Cre-KO^ cell-injected animals lacked pancreatic or metastatic lesions, in contrast to KPCY-*Mcu*^rescue^ cell-injected mice (Fig. 3D-F, S2B). To evaluate the metastatic ability of MCU-null or -expressing PDAC tumor cells, KPCY-*Mcu*^Cre-KO^ or KPCY-*Mcu*^rescue^ cells were injected into the tail veins of C57BL/6 mice. Similar to the results in the orthotopic implantation assay, KPCY-*Mcu*^Cre-KO^ tail vein-injected animals failed to form metastatic colonies, while KPCY-*Mcu*^rescue^ cells efficiently colonized the lung (Fig. 3G-I). No differences in body weight were observed (S2C). Thus, the lack of metastases seen in the orthotopic model was not solely due to the inability to form a primary lesion.

**Fig. 3.**
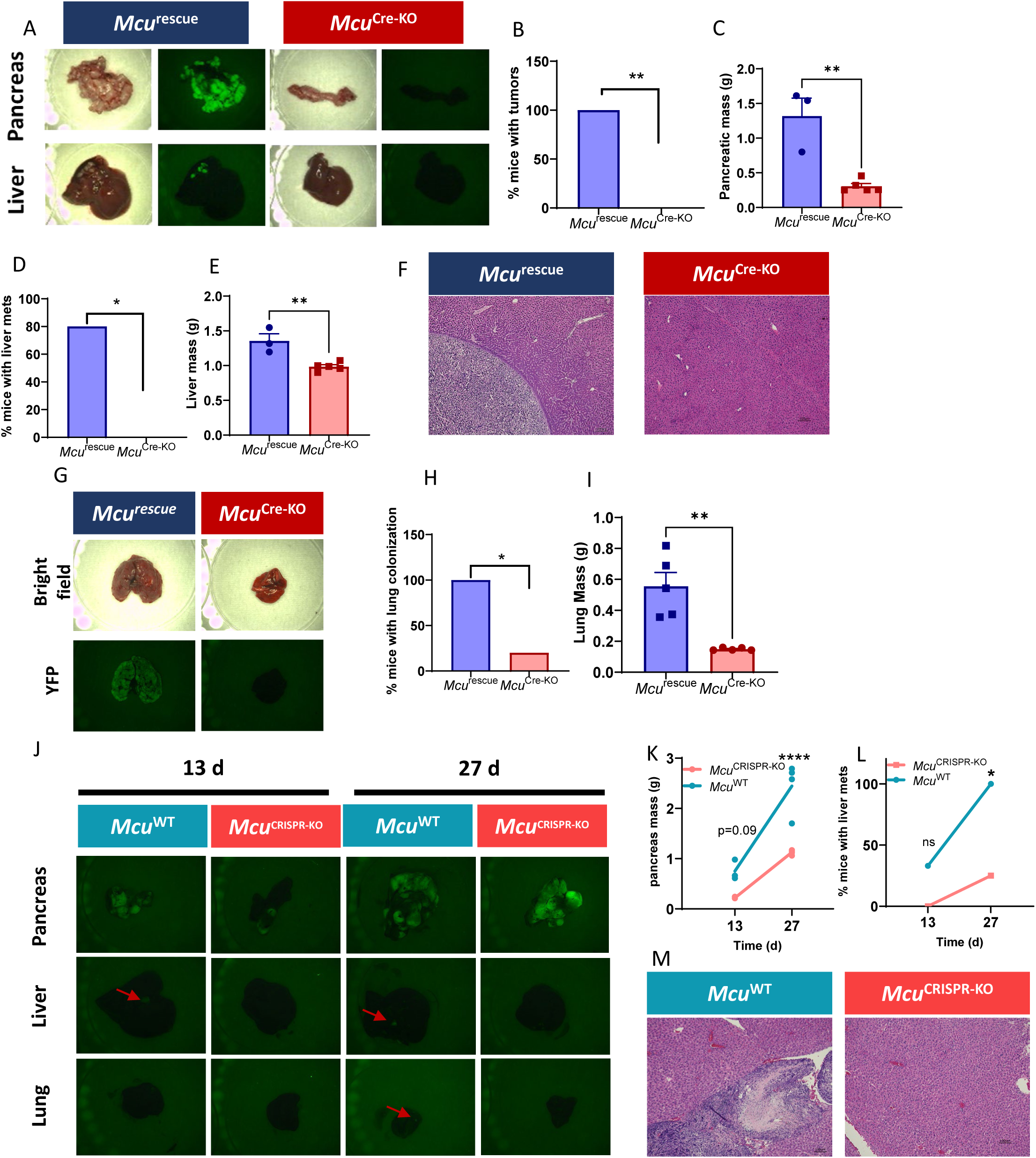
*MCU* ablation reduces tumor growth and metastasis in *in vivo* xenograft models. Mice were injected orthotopically in the pancreas with 100,000 cells (*Mcu*^Cre-KO^ or *Mcu*^rescue^) and aged for 21 days, until tumors were palpable and mice began to show symptoms. A, representative bright-field and YFP images of liver and pancreas of C57bl/6J mice injected orthotopically with *Mcu*^Cre-KO^ or *Mcu*^rescue^ cells. B, quantification of number of mice with tumors. C, total mass of pancreas. D, percent mice with liver metastases. E, liver mass of mice injected with *Mcu*^Cre-KO^ and *Mcu*^rescue^ cells. F, representative images of liver tissue stained with H&E. G, representative bright-field and YFP images of the lungs of C57/bl6J mice injected in the tail vein with 100,000 Mcu^Cre-KO^ or *Mcu*^rescue^ cells and aged for 14 days. H, quantification of lung colonization in the tail-vein injection model. I, lung mass from tail-vein injection model. J, representative bright-field and YFP images of pancreas, liver, and lung from C57/bl6 mice orthotopically implanted with 100,000 *Mcu*^CRISPR-KO^ or *Mcu*^WT^ cells in the pancreas and aged for 13 or 27 days. K, quantification of pancreatic mass. L, percent of mice with liver metastases. M, representative H&E staining of liver tissue. Number of mice with tumors or metastases were compared with Fisher’s exact test. Tissue masses were compared with Student’s t-test. *, p≤0.05. **, p≤0.01. ***, p≤0.001.

As KPCY-*Mcu*^Cre-KO^ cells failed to form tumors upon orthotopic implantation, we performed additional experiments in an orthologous model by injecting isogenic clonal cell lines generated after *in vitro* knockout of MCU (KPCY-*Mcu*^CRISPR-KO^) or control parental lines (KPCY-*Mcu*^WT^). MCU deletion reduced primary tumor burden at 13- and 27-days post-implantation (Fig. 3J-K). This parental cell line has previously been characterized as having a micrometastatic phenotype, with evidence of YFP^+^ tumor cells in the liver and lung at 27 days post-implantation (Fig. 3J, L-M and S2D). Metastatic burden was reduced in animals injected with KPCY-*Mcu*^CRISPR-KO^ cells, with little to no evidence of YFP^+^ micrometastases by fluorescent dissection microscopy or pathological analysis of H&E images (Fig. 3J, L-M and S2D). In addition, KPCY-*Mcu*^CRISPR-KO^ tumor-bearing mice had fewer ascites and spleen metastases compared with *Mcu*^CRISPR-KO^ mice (Fig. S2F). Lung lesions were not observed in *Mcu*^CRISPR-KO^ mice, whereas occasional small lung metastases (which did not affect total lung weight) were observed in *Mcu*^WT^ mice (Fig. 3J and S2E-F). Overall body weight was not affected by tumor-specific MCU deletion (S2G). Both isogenic cell lines formed poorly-differentiated tumors, with no differences in relative proportions of differentiated area (Fig. S2I). The similar results in multiple *in vivo* models of tumor cell-specific deletion of *Mcu* suggests that mitochondrial Ca^2+^ uptake significantly supports the growth and metastasis of PDAC tumors.

In the GEMM KPCY-*Mcu*^Cre-KO^ model (Fig. S1A), we did not observe improvements in survival (Fig. S1B), percent of mice with metastases (Fig. S1C), or pancreatic mass (Fig. S1D) in *Mcu*^Cre-KO^ mice compared with KPCY-*Mcu*^WT^ mice. However, *Mcu*^Cre-KO^ mice had significantly reduced liver mass (Fig. S1E), suggesting that knockout of MCU may reduce overall metastatic burden in the liver. Lung mass was not significantly different between conditions (Fig. S1F).

### MCU loss reduces EMT

We observed distinct morphological differences between MCU-KO and MCU-expressing isogenic cell lines *in vitro*. MCU-KO cells were more epithelial (semicuboidal, grew in plaque-like formations, and were less motile), whereas MCU-expressing isogenic lines had a more mesenchymal identity (spindle morphology and more motile) (Fig. 4A). We posited that these differences could be explained by mitochondrial Ca^2+^-dependent induction of an EMT program, a key pathway implicated in tumor cell plasticity and metastasis ^6,7,10,12,15,16,20,21,38^. To confirm this shift in cellular identity *in vivo*, we stained MCU-KO and MCU-expressing orthotopic tumor tissues for E-cadherin (ECAD), a marker of the epithelial state whose loss indicates EMT induction. ECAD expression was markedly reduced in MCU-expressing tumor cells (KPCY-*Mcu*^WT^ and KPCY-*Mcu*^rescue^) compared with MCU-KO cells (KPCY-*Mcu*^CRISPR-KO^ and KPCY-*Mcu*^Cre-KO^) (Fig. 4B, S3A), indicating that mitochondrial Ca^2+^ signaling facilitates EMT in PDAC tumors. Of note, KPCY-*Mcu*^WT^ cells expressed basal levels of the key EMT transcription factor, *Snai1* (Snail), whereas it was undetectable in *Mcu*^CRISPR-KO^ cells (Fig. 4C). Snail is known to repress ECAD expression and potently induce EMT downstream of TGFβ signaling, which is a canonical EMT-inducing signal in cancer^39,40^. To gain further insights into the role of mitochondrial Ca^2+^ signaling in EMT, we interrogated transcriptional differences between MCU-expressing and MCU-KO cells by RNA-sequencing. Unsupervised hierarchical clustering demonstrated that the different cell lines clustered by *Mcu* expression rather than by parental cell-line source, suggesting a strong effect of MCU expression on overall transcriptional programs (Fig. 4D, Supplemental Table 1). Consistent with the observed shift in epithelial cell identity by cell morphology and ECAD expression in MCU-deficient cells, Gene Ontology and Gene Set Enrichment Analysis (GSEA) indicated that EMT was one of the top significantly-altered gene sets between *Mcu*-expressing and *Mcu*-KO cells, with KO cells having reduced enrichment for EMT genes (Fig. 4E-F, Supplemental Tables 2-5). EMT-related genes clustered strongly by *Mcu* expression (Fig. 4G), and several EMT transcription factors were identified by CHeA3 analysis (Fig. S3B). Thus, MCU expression strongly promotes an EMT transcriptional program in PDAC.

**Fig. 4.**
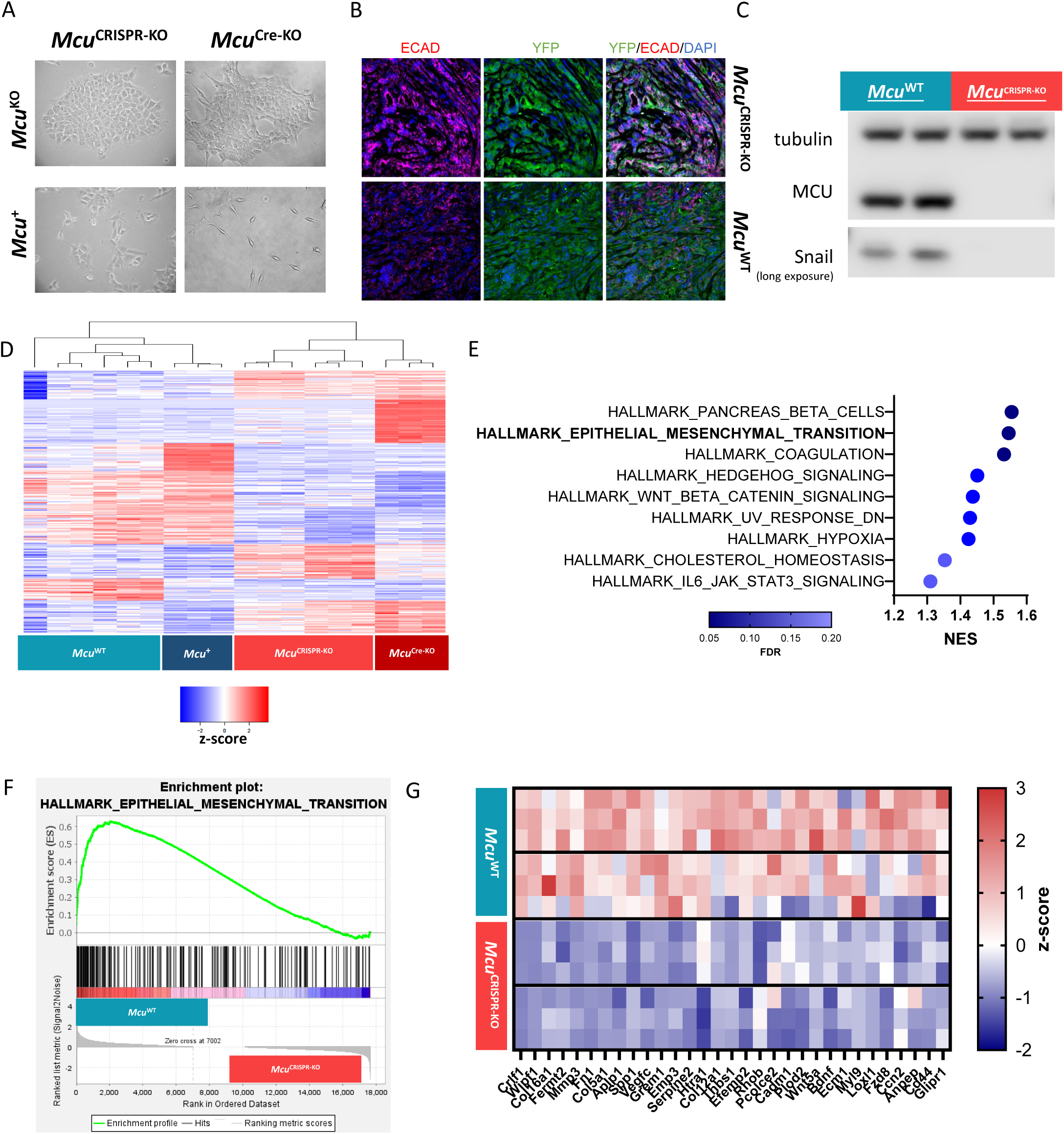
*Mcu* expression is associated with EMT. A, representative 20x bright-field images of Mcu^CRISPR-KO^, Mcu^Cre-KO^, and their isogenic control cells. B, representative immunofluorescence images of tissue from the primary lesions of 27 d *Mcu*^CRISPR-KO^ and *Mcu*^WT^ orthotopic injections from Fig. 3J-M. *Mcu*^WT^ cells express little ECAD, suggesting extensive EMT has occurred. C, western blot of Snail in *Mcu*^CRISPR-KO^ and *Mcu*^WT^ cells. D, heat map of RNAseq genes (as z-score). When unsupervised hierarchical clustering is applied to *Mcu*-knockout and *Mcu*-expressing isogenic cell lines, they independently group into *Mcu*^KO^ and *Mcu*^rescue^ groups, suggesting that *Mcu* expression strongly influences transcriptional regulation. E, GSEA enrichment pathways by normalized enrichment score (NES), colored by false discovery rate (FDR). F, GSEA enrichment analysis plot for Epithelial to Mesenchymal Transition gene set, indicating upregulation in *Mcu*^WT^ cells compared with *Mcu*^CRISPR-KO^cells. G, Heat map of top 30 leading-edge genes from GSEA of EMT genes in *Mcu*^WT^ and *Mcu*^CRISPR-KO^clones, showing clear differences between groups (results shown as z-score).

### TGFβ and Snail rescue MCU^KO^ phenotypes *in vitro*

To confirm the induction of an EMT program in MCU-expressing tumor cells, we examined protein expression levels of key EMT-pathway components. As noted, under basal, untreated conditions, *Mcu*^WT^ cells expressed the EMT transcription factor Snail at higher levels than in *Mcu*^CRISPR-KO^ cells (Fig. 4C). Higher Snail expression in *Mcu*^WT^ cells was associated with increased levels of secreted TGFβ, a known inducer of EMT, in the cell-culture media (Fig. 5A), despite similar expression of Tgfb1 transcripts (S4A-C). We speculated that the promotion of TGFβ secretion by MCU-mediated mitochondrial Ca^2+^ uptake could mechanistically link MCU expression with increased Snail expression and possibly EMT. To test the hypothesis that *Mcu*^CRISPR-KO^ cells expressed lower levels of EMT markers as a consequence of lower secretion of EMT-induction factors, we used TGFβ treatment and stable *Snai1* overexpression (Snail^OE^), both well-characterized orthogonal methods of EMT induction, in the isogenic *Mcu*^WT^ and *Mcu*^CRISPR-KO^ cell lines. Treatment with the EMT-inducing ligand TGFβ (10 ng/ml for 72h) reduced the epithelial cell marker ECAD and increased the expression of mesenchymal markers N-cadherin (NCAD), Vimentin, and Snail, independent of *Mcu* status (Fig. 5B). Of note, *Mcu*^CRISPR-KO^ cells expressed higher levels of the TGFβ receptor (*Tgfbr1)* transcripts and similar levels of *Tgfbr2* (Fig. S4E) compared with *Mcu*^WT^ cells. Furthermore, similar ΕΜΤ phenotypes were induced in both cell lines by stable *Snai1* overexpression (Fig. 5C-D). Thus, functional EMT machinery remains intact in *Mcu*^CRISPR-KO^ cells, despite their lower levels of basal Snail expression and absence of EMT phenotypes.

**Fig. 5.**
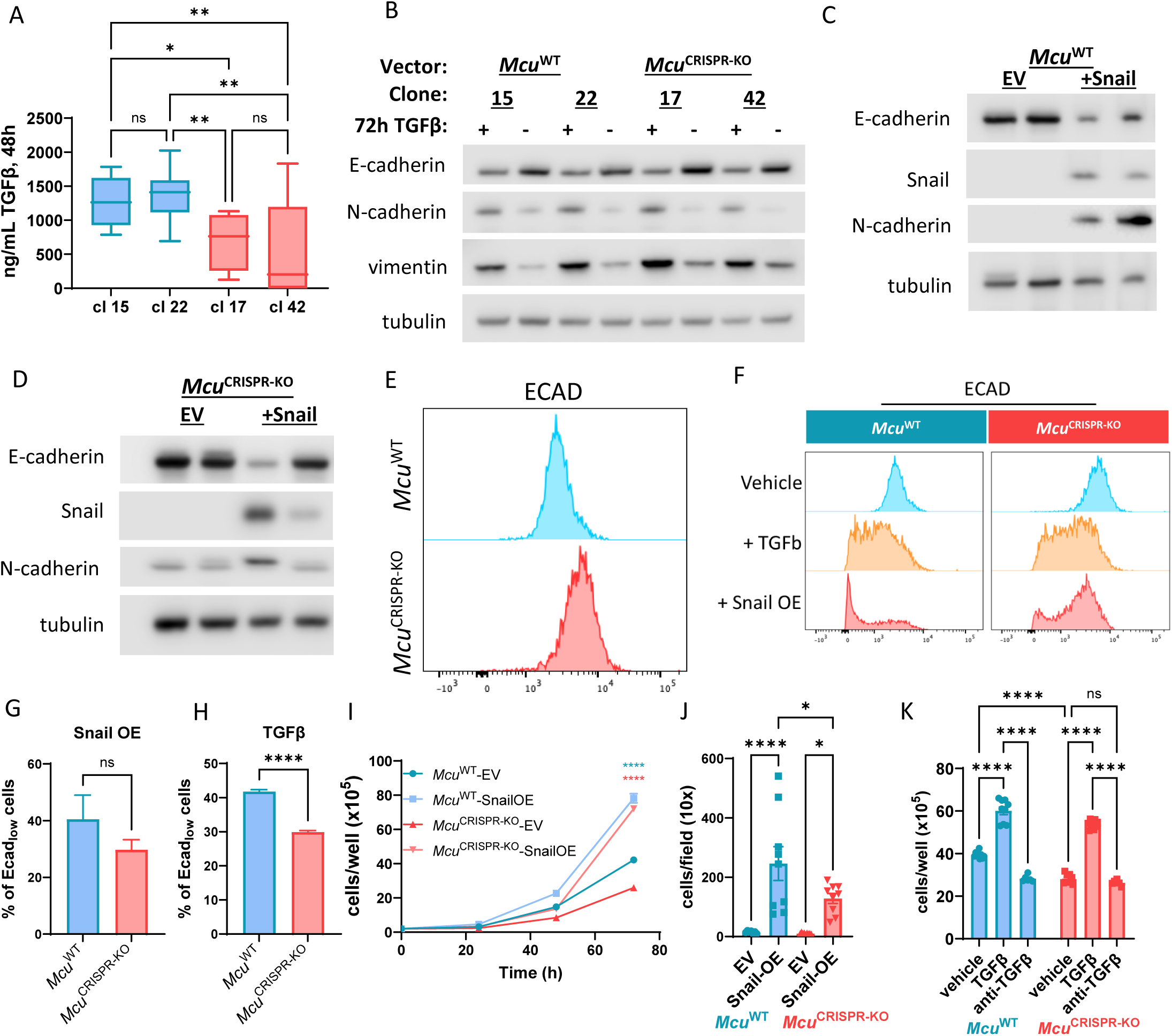
Exogenously induced EMT ameliorates many deficits seen in *Mcu*^CRISPR-KO^ cells. A, *Mcu*^WT^ cells secrete more TGFβ into the media, as measured by ELISA of the media at 48 h. B, Exogenous treatment with 10 ng/mL TGFβ for 72 h increases N-cadherin and vimentin protein levels and reduces E-cadherin levels independent of MCU expression. C, D, Stable *Snai1* expression increases Snail and N-cadherin expression and reduces E-cadherin levels in *Mcu*^WT^ cells (C) and *Mcu*^CRISPR-KO^ cells (D). E, Flow cytometry plot for ECAD surface expression of untreated *Mcu*^WT^ and *Mcu*^CRISPR-KO^ cells indicates that *Mcu*^CRISPR-KO^ cells express more ECAD. F, flow cytometry plots for *Mcu*^CRISPR-KO^ and *Mcu*^WT^ cells treated with 10 ng/mL TGFβ or stable Snail expression (quantified in G and H, respectively). I, Snail expression increases cell growth of *Mcu*^CRISPR-KO^ cells to levels comparable to that of *Mcu*^WT^ cells. J, Snail overexpression increases 24 h transwell migration of *Mcu*^CRISPR-KO^ cells to a level intermediate between *Mcu*^WT^ cells with or without Snail overexpression. K, 72 h cell counts indicate that TGFβ increases *Mcu*^CRISPR-KO^ cell growth to levels comparable to *Mcu*^WT^ cells treated with TGFβ. Treatment with a TGFβ neutralizing antibody (anti-TGFβ) significantly reduced proliferation in *Mcu*^WT^, but not *Mcu*^CRISPR-KO^ cells. Cell count data are analyzed with two-way ANOVA with Tukey’s posthoc. Two group data are analyzed with Student’s t-test. *, p≤0.05. **, p≤0.01. ***, p≤0.001.

Western blot analysis of whole-cell lysates, which measures total ECAD protein expression throughout the cell, gives an incomplete picture of epithelial versus mesenchymal identity. Rather, membranous ECAD (M-ECAD) is important for maintaining epithelial cell identity and loss of ECAD from the membrane is associated with the EMT phenotype ^6,12,15,20,21^. Flow cytometric quantification of surface ECAD in non-permeabilized cells revealed that *Mcu*^CRISPR-KO^ cells have higher baseline surface ECAD expression than *Mcu*^WT^ cells (Fig. 5E), consistent with their more epithelial nature. Induction of EMT by overexpression of *Snai1* or TGFβ treatment more strongly reduced surface ECAD levels in *Mcu*^WT^ cells relative to *Mcu*^CRISPR-KO^ cells (Fig. 5F-H). Collectively, these results indicate that MCU-expressing cells are poised to undergo EMT during tumorigenesis through enhanced expression of EMT transcription factors, and that inhibition of mitochondrial Ca^2+^ uptake abrogates the ability of pancreatic tumor cells to lose their epithelial cell identity, which has important implications for their metastatic ability.

Remarkably, *Snai1* overexpression rescued *Mcu*^KO^-associated deficits in tumor cell clonogenicity (Fig. S4E), proliferation (Fig. 5I), wound healing (Fig. S4F) and transwell migration (Fig. 5J) to levels comparable to those of *Mcu*^WT^ cells. TGFβ treatment also increased cell proliferation in both groups (Fig. 5K). TGFβ neutralizing antibody reduced unstimulated growth of *Mcu*^WT^ cells, but had no effect on basal proliferation of *Mcu*^CRISPR-KO^ cells (Fig. 5K), consistent with higher TGFβ production in *Mcu*^WT^ cells (Fig. 5A) and indicative of divergent TGFβ signaling upon loss of *Mcu*. Similarly, wound healing was also increased by TGFβ treatment in both *Mcu*^CRISPR-KO^ and *Mcu*^WT^ cells, and treatment with the TGFβ neutralizing antibody reduced this phenotype only in the *Mcu*^WT^ cells (Fig. S4G). In the *Mcu*^Cre-KO^ and *Mcu*^rescue^ models, TGFβ increased proliferation only of the MCU-expressing cells (Fig. S4H). Overall, these results suggest that deletion of inhibition of mitochondrial Ca^2+^ uptake reduces malignant phenotypes, in part, through reducing cell-autologous secretion of pro-tumorigenic signals, but loss of MCU does not prevent the responses to these signals. Further, EMT induced by stable Snail expression or TGFβ can induce key malignant phenotypes in tumor cells lacking MCU-mediated mitochondrial Ca^2+^ signaling, highlighting molecular redundancy in the EMT pathway.

### Snail expression rescues *Mcu*^CRISPR-KO^ phenotypes *in vivo*

To observe if the phenotypic changes observed *in vitro* upon Snail overexpression were maintained *in vivo*, we implanted *Mcu*^CRISPR-KO^ cells stably expressing Snail or empty vector (EV) into the pancreas of C57BL/6 syngeneic mice. Snail expression increased the primary tumor burden (Fig. 6A-B) and enhanced their metastatic ability (Fig. 6C). Increased tumor and metastatic burden were associated with decreased surface ECAD expression in YFP^+^ tumor cells (Fig. 6D), indicative of tumor cells undergoing EMT and entering into the metastatic cascade. Continued expression of ECAD in adjacent YFP-negative wild-type ductal cells in Snail^OE^ tumors (Fig. 6D) indicates that cell-intrinsic mechanisms within tumor cells mediate EMT induction in this context. The loss of ECAD expression and appearance of more mesenchymal morphology in histological sections strongly indicates that overexpression of Snail robustly induces EMT in *Mcu*^CRISPR-KO^ cells *in vivo.* Therefore, inhibition of mitochondrial Ca^2+^ signaling appears to reduce the propensity of cells to cell-autologously induce EMT, but not the ability of cells to respond to EMT induction from exogenous sources. These findings have implications for targeting of MCU.

**Fig. 6.**
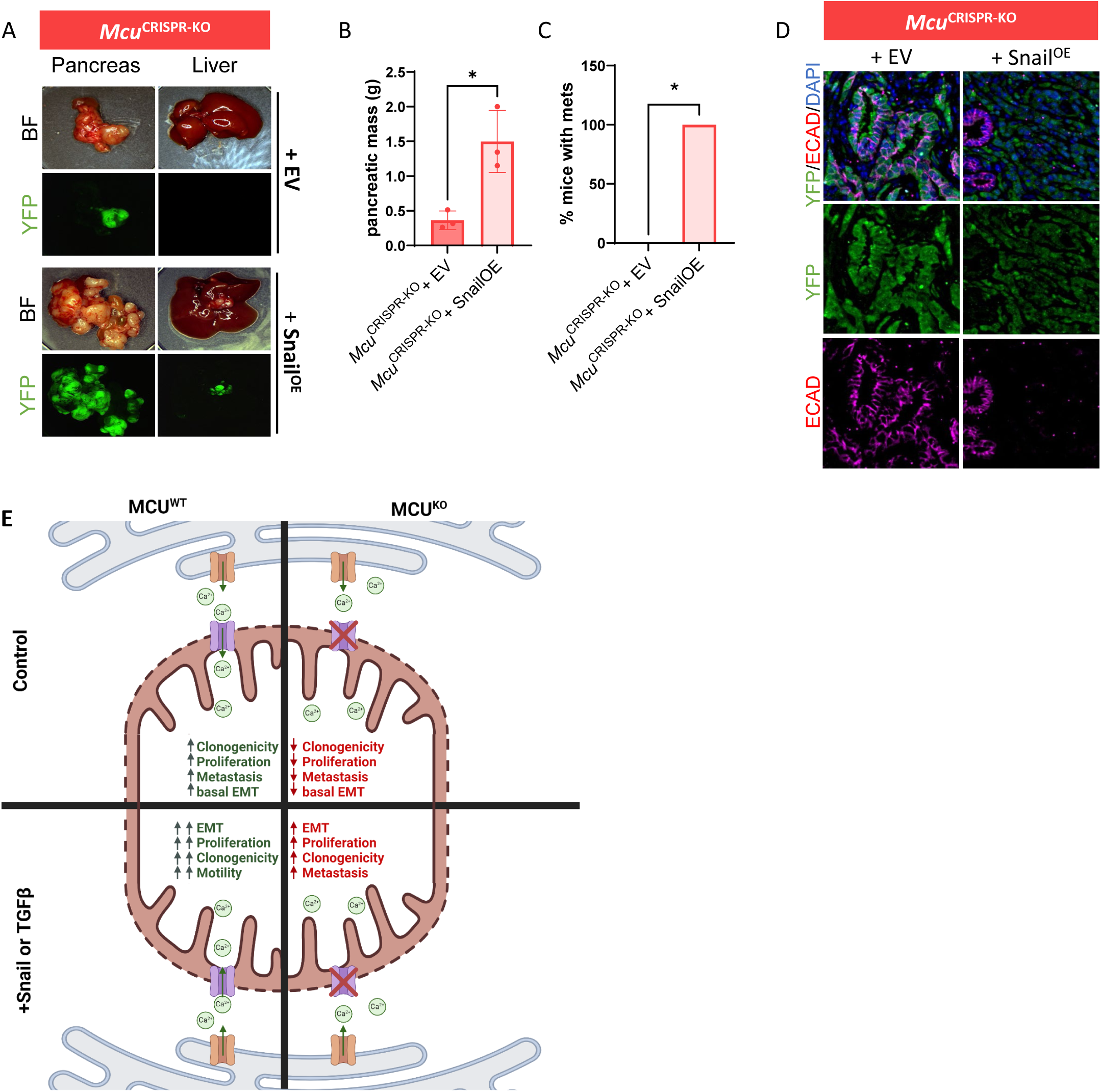
A, representative bright field (BF) and YFP images of pancreas and liver from C57/bl6J mice orthotopically implanted with 100,000 *Mcu*^CRISPR-KO^+EV or *Mcu*^CRISPR-KO^+*Snail*^OE^ cells for 21 days. B, pancreatic mass from orthotopic injection model. C, percent of mice with metastases. B, representative immunofluorescence images of *Mcu*^CRISPR-KO^+EV or Mcu^CRISPR-KO^+*Snail*^OE^ cells stained for YFP (lineage tracer), DAPI (nuclear marker), and ECAD (epithelial marker). *Mcu*^CRISPR-KO^+EV cells show robust staining of ECAD in YFP-expressing tumor cells, but YFP^+^ *Mcu*^CRISPR-KO^+*Snail*^OE^ cells only poorly co-express ECAD. A resident, normal-type duct is shown to demonstrate that epithelial cells in the host robustly express ECAD. E, schematic of effects of MCU deletion in the presence and absence of exogenous EMT induction with stable Snail overexpression or TGFβ treatment. Two group data are analyzed with Student’s t-test, and proportion data is analyzed by Chi square. *, p≤0.05. **, p≤0.01. ***, p≤0.001.

## Discussion

Here, we identify mechanisms by which the mitochondrial Ca^2+^ influx channel MCU supports oncogenic and pro-metastatic functions of pancreatic tumor cells in murine KPCY models of PDAC. MCU deletion reduces PDAC cell motility, clonogenicity, and proliferation *in vitro* and tumor growth and metastasis *in vivo*. Mechanistically, inhibition of mitochondrial Ca^2+^ uptake restricts tumor cell plasticity by reducing EMT and entry into the metastatic cascade, ultimately blocking their ability to colonize distant niches.

Mitochondrial Ca^2+^ homeostasis plays dual roles in cells. Excessive mitochondrial uptake can result in Ca^2+^ overload and cell death by apoptotic and necrotic mechanisms. Conversely, mitochondrial Ca^2+^ is a critical control mechanism for the regulation of basal bioenergetics and for enhanced energy production during periods of increased metabolic demands that are likely encountered during multiple steps of the metastatic cascade. Cancer cells may have a “Goldilocks zone” for Ca^2+^ signaling through MCU, wherein sufficient influx is necessary to support proliferation and metabolism, but excessive signaling contributes to toxicity. This set-point may differ significantly based on cell type, environmental conditions, driver mutations, or expression levels of other proteins within the signaling pathway. Such differences could contribute to observed variability in the effects of MCU expression on survival and tumor growth in different cancers. For example, according to TCGA datasets, high MCU-expression levels in melanoma and kidney tumor tissues are associated with enhanced patient survival, in contrast to the opposite associations observed in PDAC, liver, and breast cancer patients^41^. We previously showed that inhibition of ER-to-mitochondrial Ca^2+^ flux is selectively toxic for cancer cell lines, which may suggest that its inhibition could be tolerated by patients while maintaining anticancer efficacy. However, only a few, non-selective agents that directly modulate MCU activity have been identified^42,43^. In the absence of reliable means to pharmacologically inhibit MCU, we have demonstrated through proof-of-principal genetic deletion experiments that MCU drives PDAC disease aggressiveness and is an attractive target for future studies. Additionally, whole body knockout of MCU in outbred mice has few effects ^25,29,44^, highlighting the potential of targeting MCU as a therapeutic vulnerability in cancer.

We found that lack of MCU expression in PDAC restricts EMT. However, when exogenous EMT-inducing pressures were applied through stable expression of Snail or application of exogenous TGFβ, *Mcu*^KO^ cells remained competent to undergo EMT. Such plasticity suggests that extrinsic induction of EMT, for example by secretion of factors like TGFβ by cancer-associated fibroblasts, may reduce the effectiveness of MCU inhibition as an anticancer target. Such findings may at least partially explain the lack of a phenotype in our genetic model of MCU-deletion in the KPCY background, in contrast to the striking phenotypes in the xenograft models. While these observations may suggest that targeting MCU as a mono-therapeutic approach might not be fruitful, it is possible that a combinatorial approach, for example with inhibitors of TGFβ signaling ^45–47^. Such approaches emphasize the need to develop specific pharmacology for MCU-mediated mitochondrial Ca^2+^ uptake that is currently lacking.

While the reduction of malignant phenotypes and basal tendency toward EMT in the context of MCU deletion is at least partially due to inhibition of cell-autologous secretion of pro-tumorigenic factors such as TGFβ, other processes likely contribute as well. These may include alterations in metabolism, activation states of other signaling pathways, or even proclivity for senescence. Notably, EMT phenotypes have been linked to metabolic alterations, and mitochondrial Ca^2+^ is known to be an important regulator of ATP production and the synthesis of biochemical intermediates produced by flux through the TCA cycle^6,12,18,19^. Crucially, cancer-associated fibroblasts may be able to help support MCU^KO^ cells in stromally-rich GEMM models by secreting metabolic intermediates and growth factors to support tumor growth despite impairments in the absence of functional mitochondrial Ca^2+^ signaling through the MCU complex. Future studies to examine the role of these secreted factors in GEMM and xenograft models of PDAC in the context of MCU in both basal and EMT-induced contexts are thus warranted. A deeper understanding of the relationships among tumor cells and supportive stromal cells will further inform our understanding of the potential of targeting MCU in PDAC.

## Methods

### Reagents and cell lines

Panc-1 (CRL-1469) and MiaPaCa2 (CRL-1420) cells were obtained from ATCC. All murine parental cell lines were developed from experimental mice as previously described ^37^. HPDE (Kerafast H6c7) were grown in Keratinocyte SFM + EGF + bovine pituitary extract (Invitrogen 17005042) supplemented with 1x antibiotic-antimycotic (A/A, Gibco 15240-062). All other cell lines were maintained in DMEM (Corning 10-013CM) + 10% fetal bovine serum (FBS, Hyclone SH30071.03) + 1x A/A. All cell lines were maintained in a humidified incubator at 37°C and 5% CO_2_ with media changes or passaging every 2-3 days.

For all stable clones, single-cell clones were developed by transfection with Lipofectamine 3000 (Thermo Fisher Scientific, L3000001) for 48 h, then selection of a polyclonal cell line with a given antibiotic and subsequent isolation of single-cell clones via limiting dilution. When possible, control lines expressing empty vectors were used as negative controls. Clones were verified for MCU expression by western blot (WB) and mitochondrial Ca^2+^ uptake assay. KPCY-*Mcu*^rescue^ lines were generated by stable expression of pCMV-*Mcu*-V5-His-puro, selected with 8 µg/mL puromycin then maintained under 2 µg/mL puromycin. For KPCY-*Mcu*^CRISPR-KO^, pLenti-CRISPR-V2-sgMCU-mCherry (a generous gift from Mohamed Trebak) transfected cells were selected by limiting dilution. For Snail expression, KPCY-*Mcu*^CRISPR-KO^ and isogenic KPCY-*Mcu*^WT^ cells were transfected with pCDH-Snail-puro and selected with 8 µg/mL puromycin, then maintained at 2 µg/mL thereafter. When indicated, cells were treated with 10 ng/mL TGFβ (Millipore Sigma SRP3171) in culture; this was replenished every 2 days.

### Cell proliferation assays

20,000 cells/well were plated in 24-well tissue culture treated plates in 2 mL of media unless otherwise noted. At given time points, media was aspirated, cells were rinsed with 1 mL 1x DPBS, and wells were trypsinized with 250 μL 0.25% trypsin for ∼3 min until detachment. Trypsinized cells were -mixed with 250 μL complete, FBS-containing media and counted manually by hemocytometer, with the average of two technical replicates taken as the value-. Three separate wells were counted at each time point per condition, and three independent cell count experiments were carried out.

### Pancreatic cancer patient samples

Human PDAC or normal pancreatic tissues were obtained from the UMass Center of Clinical and Translational Sciences Biorepository and derived retrospectively from patients undergoing surgery at UMass Memorial Hospital consented under the IRB approved protocol no. H-4721. De-identified FFPE tumor specimens were cut into 5μm sections and IHC staining was performed as described above. Briefly, MCU primary antibody was stained at 1:200 (Sigma #HPA016480).

### Tissue staining and imaging

Tissues were isolated from mice, placed in cassettes in zinc formalin fixative, and stored at 4°C overnight. Then, tissue cassettes were transferred to 70% ethanol in distilled water at 4°C until further processing. All tissues were paraffin embedded, sectioned, and stained with hematoxylin and eosin by the Molecular Pathology and Imaging Core (MPIC) at the University of Pennsylvania (Center for Molecular Studies in Digestive and Liver Diseases - P30DK050306), RRID: SCR_022420).

For immunofluorescent staining, tissue sections were deparaffinized in Xylene, rehydrated, and antigen retrieval was performed with R-Buffer A (Electron Microscopy Sciences 62706-10). Slides were blocked and permeabilized for 1 h at room temperature with 5% donkey serum in 0.3% PBS-Triton X and then slides were left in primary antibody in 5% donkey serum in 0.3% PBS-Triton X overnight at 4°C. Primary antibodies used were MCU (HPA016480, Sigma-Aldrich), ECADHERIN (Clone M108, Takara), Ki-67 (ab16667, Abcam), and green fluorescent protein (ab6673, Abcam), which recognizes YFP. Slides were mounted in Fluoromount-G™ Mounting Medium with DAPI (Invitrogen 00-4959-52). Imaging was completed on a Leica Thunder Tissue Imager and analyzed on QuPath^48^.

### Invasion and migration assays

Cultured cells were plated at a density of 20,000 cells/well in the top of transwell invasion (Millipore Sigma ECM550, following manufacturer’s instructions) or migration (Corning, 3464) plates in serum-free DMEM. Complete DMEM + 10% FBS was used as an attractant in the bottom of the plate. After 24 h, the tops of the wells were cleaned and cells on the bottom of the membrane were fixed in 4% paraformaldehyde (Electron Microscopy Sciences, 15713) and stained with 10 μg/mL DAPI in 1% Triton X100 (Sigma T9284). Three different fields of view were imaged at 20x and analyzed by ImageJ.

### Wound healing assays

Cultured cells were plated in complete media at 500,000 cells/well of 12-well tissue culture treated plates and incubated overnight. The next day, monolayers were scratched by hand with a 200 µL pipet tip to create a wound. Media was changed, and plates were imaged at 10x. Plates were incubated for 18-24 h as noted then imaged again in the same conditions, unless otherwise noted. Relative migration area was calculated by: 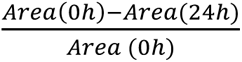 were repeated in triplicate, with biological replicates in triplicate per experiment.

### Clonogenic assays

Cells were plated at 10,000 cells/well in 6-well tissue culture treated plates and incubated for 7 d. Cells were then rinsed with 1x DPBS, incubated at room temperature in crystal violet fix/stain solution (1% methanol, 1% paraformaldehyde, 0.5% crystal violet in DPBS) for 1 h, then gently rinsed with water until the wells run clear. After drying overnight, the plates were imaged with a GeneSys GBox and quantified for clonogenic area with ImageJ using the ColonyArea plug-in.

### Spheroid formation assays

Tumorsphere formation assays were carried out as previously described with a few changes ^49^. Briefly, 200 cells were plated in 200 μL media in low-adhesion plates coated with Aggrewell Antiadherence solution (Stem Cell Tech #07010) according to manufacturer instructions. Outer wells were filled with 1xDPBS to reduce evaporation. Spheroids were counted manually on a microscope after 7 d of incubation.

### Western blotting

Media was aspirated from culture plates and cells were rinsed with 1x DPBS before thorough aspiration. Cells were then harvested in RIPA buffer + 200 μM PMSF + 1x cOmplete Mini protease inhibitor cocktail (EDTA free, Roche 11836170001) via scraping on ice, transferred to labeled tubes, and rotated for 1.5-2 h at 4°C. Samples were then centrifuged at 13000+ rpm and 4°C for 10 min and transferred to new tubes and quantified via Pierce BCA assay according to manufacturer’s instructions, with samples diluted 1:5. Samples were mixed to load 10-20 μg protein/well with 4x Laemmli buffer (Biorad 1610747) according to manufacturer specifications. Precision Plus Dual Color Standard (Biorad 161-0374) served as protein Standard. Samples were run on NuPAGE 4-12% bis/tris mini protein gels (NP0321-0323) at 100 V in MES running buffer (NP0002) for ∼1 h then transferred for 1 h at 100 V onto Immobilon-P PVDF membrane on ice. After blocking for 1.5 h at room temperature on a rocker in 5% w/v dry milk in TBST (1x tris buffered saline + 0.1% Tween 20), blots were incubated overnight on a shaker at 4°C. After rinsing, blots were incubated in secondary antibody (Cell Signaling Technologies: anti-mouse-HRP, 7076, and/or anti-rabbit-HRP, 7074). Blots were then rinsed with TBST and visualized with a GeneSys GBox and SuperSignal West PICO Plus ECL reagent (ThermoFisher Scientific, 34577). Results were quantified as appropriate with ImageStudio Lite.

### Murine studies

Orthotopic implantation of tumor cells was performed as previously described ^50,51^. Briefly, mice were anesthetized with isoflurane and a sterile field around the abdomen was prepared. An incision was made in the upper left quadrant of the abdomen and the body of the pancreas was exposed. Then 1.0x10^5^ cells in 100 μL sterile DMEM were injected into the tail of the pancreas with an insulin syringe. KPCY-derived tumor cells from C57BL/6J mice were injected into C57BL/6J mice (000664, The Jackson Laboratory). The formation of a liquid bleb at the injection site verified a successful injection. After injection, a sterile pad was held to the injection site to prevent tumor cells from leaking into the abdominal cavity. Finally, the pancreas was placed back into the abdomen and then the peritoneum and skin were sutured closed with 4-0 coated sutures.

For tail vein injections, an insulin syringe was loaded with 1.0x10^5^ KPCY cells in 100 μl sterile DMEM and injected into the tail vein of C57BL/6 animals (000664, The Jackson Laboratory). Lungs were harvested at the indicated time points, imaged for YFP and bright field, and then formalin fixed for downstream analysis.

### Flow cytometry

Surface ECAD levels were measured by a previously described method [PMID: 34324787]. Briefly, cells were removed from culture plates and dissociated into single cells using Hank’s Enzyme Free Cell Dissociation Solution (S-004-C, EMD Millipore). Cells were stained using anti-ECAD (147308, BioLegend) or isotype control (400418, BioLegend) in FACS buffer for 15 min on ice in the dark. Cells were washed in FACS buffer and then stained with DAPI and filtered through a 70-μm strainer to create a single cell suspension. Flow cytometry was run on an LSR II at the University of Pennsylvania Flow Cytometry Core. These experiments measure surface ECAD levels only, as cells were not permeabilized.

### RNASeq

RNA was isolated with a RNeasy Mini kit (Qiagen 74104) from 10-15-cm dishes and sequenced by Novogene with a NovoSeq PE150 at ∼20 M paired-end reads. Raw reads were processed with Salmon and DESeq2 before analysis with GSEA and GO. At least 3 biological replicates were used for each experimental condition, and principal components were used to verify the reproducibility of replicates.

### Statistics

Unless otherwise noted, all experiments were carried out as 3 separate, independent experiments with at least 3 biological replicates per experiment. Data were analyzed with GraphPad Prism (versions 8-10) or R, unless otherwise noted. For all normally-distributed two-group data, Student’s T-test was used. For multigroup, one-independent variable data, one-way ANOVA with Sidak’s posthoc was used. When two independent variables were present (i.e., in ±MCU, ±Snail experiments and time courses), two-way ANOVA with Sidak’s posthoc was employed. All data were assessed for normality with Kolmogorov Smirnov tests and for outliers with ROUT with Q=5% before analysis.

### Conflicts of Interest

B.Z.S. receives research funding from Boehringer-Ingelheim and Revolution Medicines and holds equity in iTeos Therapeutics. J.R.P. receives research funding from Boehringer-Ingelheim. This research was supported by the Penn Pancreatic Cancer Research Center.

## Acknowledgements

We thank the Penn Metabolomics Core (RRID:SCR_022381) in the Cardiovascular Institute at the University of Pennsylvania for metabolomics analyses and the Molecular Pathology and Imaging Core (MPIC) at the University of Pennsylvania (Center for Molecular Studies in Digestive and Liver Diseases - P30DK050306), RRID: SCR_022420). In addition, we thank J. Dylan Weissenkampen for his assistance with bioinformatics. This work was supported by an American Gastroenterology Association Bern Schwartz Research Scholar Award in Pancreatic Cancer (J.R.P.); NIH/NCI K99-R00 CA252153 (J.R.P.); NIH/NCI R01CA250173 (J.K.F.), NIH/NCI F32-CA250144 (J.S.W.), and the Penn Pancreatic Cancer Research Center. C.J. is supported by the Initiative for Maximizing Student Development (IMSD) T32 at UMass Chan Medical School (T32 GM135751). J.P. is supported by the Innate Immunity Training Program (IITP) T32 at UMass Chan Medical School (T32 AI095213).

## Supplemental Figure Legends

**Fig. S1.**
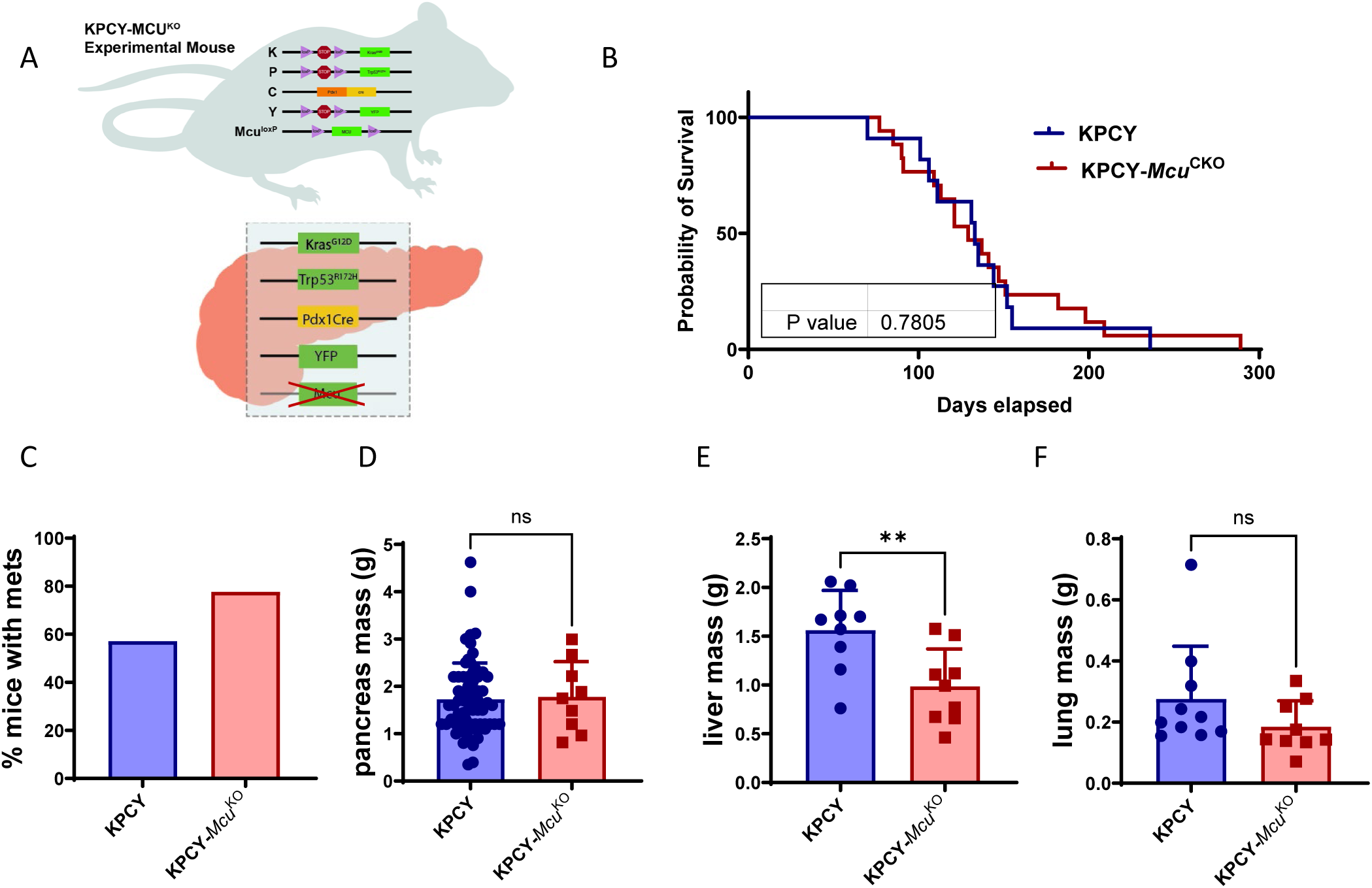
*Mcu*^KO^ does not affect overall survival in the KPCY GEMM model of PDAC. A, ***<u>K</u>****ras^G12D^; T**<u>p</u>**53^R172H^; Pdx1-**<u>C</u>**re; R26^LSL-Yfp/LSL-**Y**fp^* (i.e. ‘KPCY’) mice were bred with or without *Mcu*^fl/fl^ alleles to generate GEMMs. B, overall survival was not altered in KPCY-*Mcu*^Cre-KO^ mice vs. KPCY-*Mcu*^WT^ controls by Kaplan Meier curve. C, percent of mice with metastases per group were not significantly different. D, pancreatic mass of KPCY-*Mcu*^WT^ and KPCY-*Mcu*^Cre-KO^ mice. E, liver mass. F, lung mass.

**Fig. S2.**
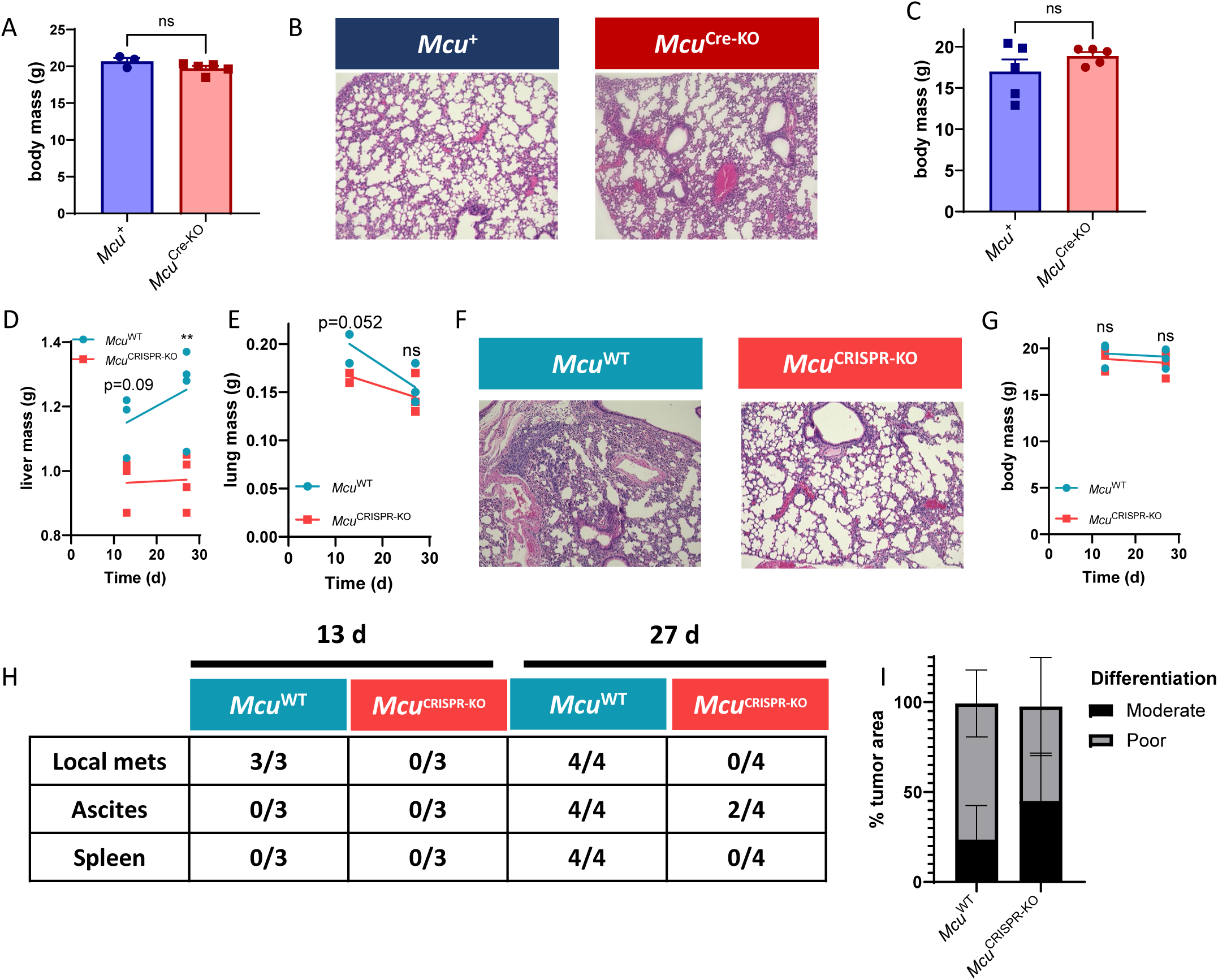
MCU ablation reduces tumor growth and metastasis in *in vivo* xenograft models. A, total body mass was not altered in mice orthotopically injected with *Mcu*^Cre-KO^ vs *Mcu*^rescue^ cells in the pancreas. B, Representative lung images, showing no malignant infiltration. C, Body mass of mice injected with KPCY-*Mcu*^Cre-KO^ or KPCY-*Mcu*^rescue^ cells in the tail vein. Liver mass (D), lung mass (E), representative lung images (F), body mass (G), quantification table of local metastasis (H), and percent differentiation of primary lesions (I) in a 13 and 27 day pancreatic orthotopic xenograft model. All two-group data analyzed by Student’s t-test. *Mcu*^CRISPR-KO^ data is analyzed with two-way ANOVA.

**Fig. S3.**
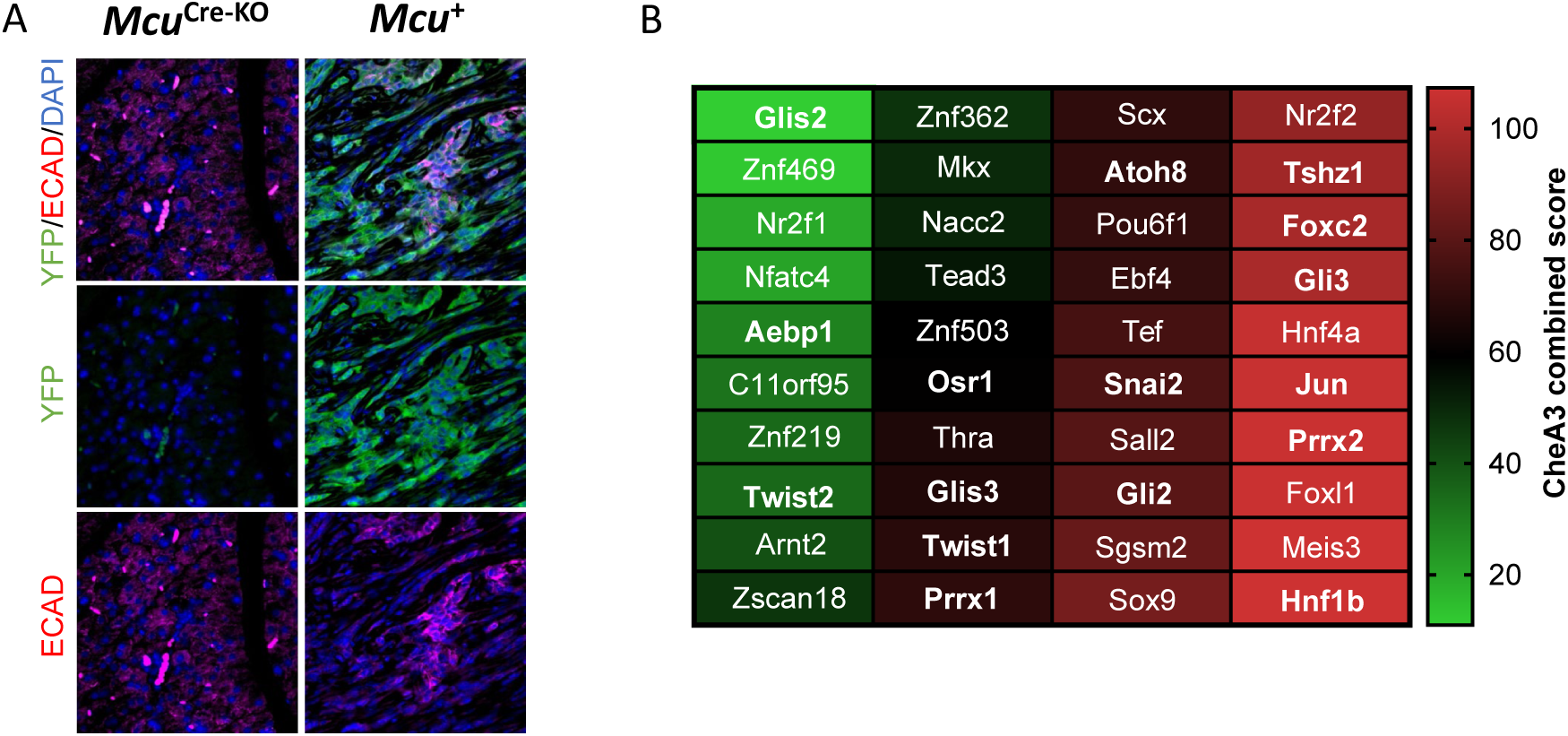
MCU expression is associated with epithelial to mesenchymal transition. A, representative immunofluorescence images of *Mcu*^Cre-KO^ and *Mcu*^rescue^ tumor tissue shows that YFP+ *Mcu*^Cre-KO^ cells are more epithelial than *Mcu*^rescue^ cells. B, CheA3 combined scores for *Mcu* expression. EMT-related transcription factors in bold.

**Fig. S4.**
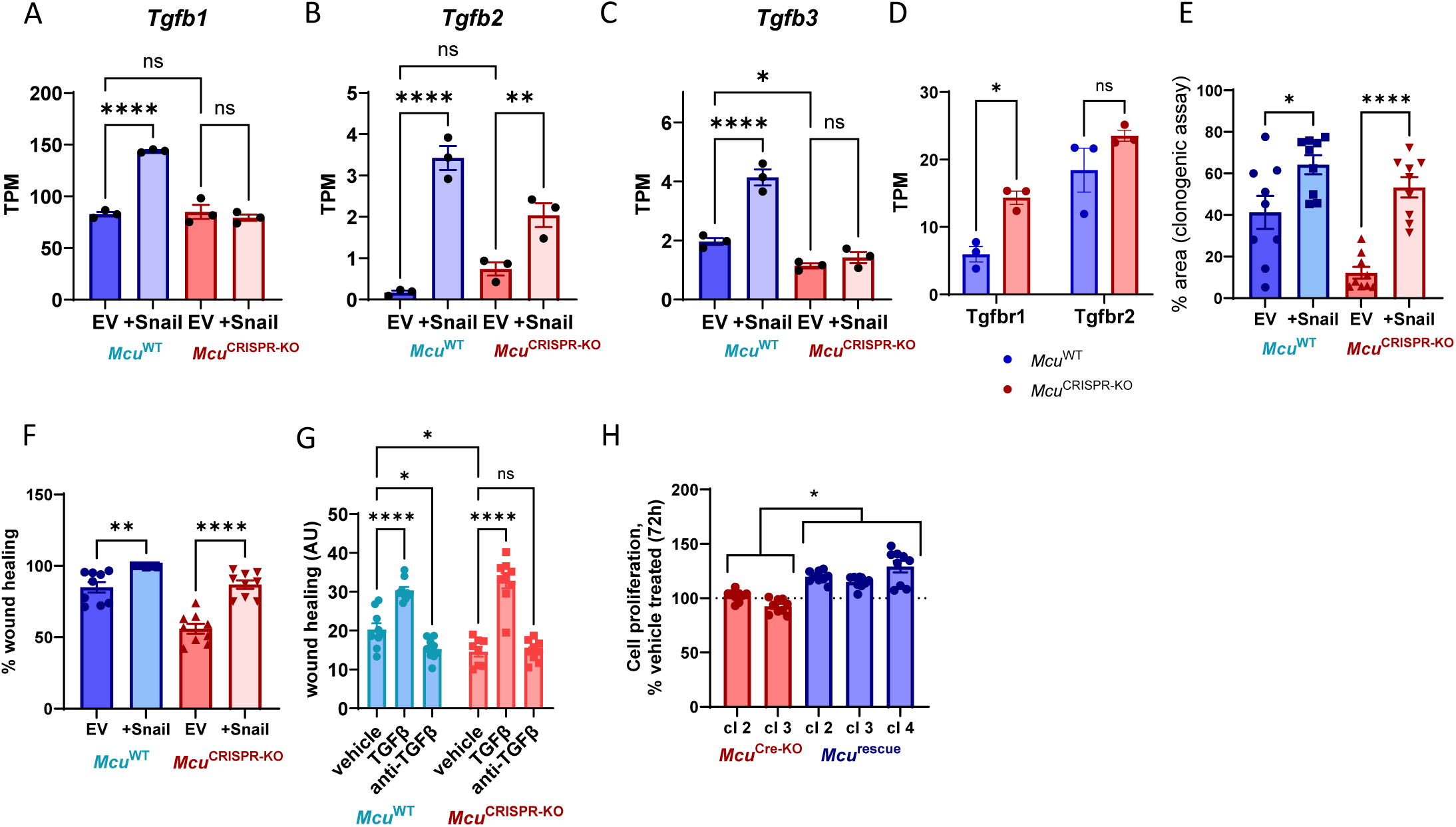
Exogenously induced EMT ameliorates many deficits seen in *Mcu*^CRISPR-KO^ cells. Transcriptional abundance of *Tgfb1* (A), *Tgfb2* (B), and *Tgfb3* (C) are generally similar in *Mcu*^WT^ and *Mcu*^CRISPR-KO^ cells +EV. Snail expression increases levels of all these transcripts in the context of *Mcu*^WT^, but only a slight increase in *Tgfb2* is seen in the context of *Mcu*^CRISPR-KO^. D, transcriptional abundance of *Tgfb1* and *Tgfb2* transcripts are increased in *Mcu*^CRISPR-KO^ cells. E, 7 d clonogenic assay of KPCY-*Mcu*^CRISPR-KO^ and *Mcu*^WT^ ± stable Snail expression. F, 24h wound healing assay of KPCY-*Mcu*^CRISPR-KO^ and *Mcu*^WT^ ± stable Snail expression. G, KPCY-*Mcu*^Cre-KO^ and KPCY-*Mcu*^rescue^ cells were treated with TGFβ for 72 h, then counted and normalized to vehicle treated cells. All data analyzed by one-way ANOVA with Sidak’s posthoc. H, *Mcu*^Cre-KO^ cells did not proliferate more quickly when treated with 20 ng/mL TGFβ for 72 h, but *Mcu*^rescue^ cells had ∼25% more proliferation under this treatment paradigm. *, p≤0.05. **, p≤0.01. ***, p≤0.001.

## Supplemental Table Legends

**Supplemental Table 1. Gene counts (in transcripts per million, TPM) for Cre mediated (1151) and CRISPR-mediated *Mcu*^KO^ (2838) cell lines. Each cell line was assayed in triplicate.** 1151MCU, *Mcu*^rescue^ cells. 1151EV, *Mcu*^Cre-KO^. 2838EV15 and 2838EV22, two separate clones of *Mcu*^WT^. 2838sgMCU17 and 2838sgMCU42, two separate clones of *Mcu*^CRISPR-KO^.

**Supplemental Table 2. GO pathway analysis for Biological Process, Cellular Component, Molecular Function, and Reactome Pathways for pathways which are significantly upregulated in *Mcu*^WT^ 2838 clones over *Mcu*^CRISPR-KO^.** All genes significantly upregulated in *Mcu*^WT^ (p. adj.<0.05) were used for GO analysis. REFLIST, total number of genes in given annotation data set. GENESET, total number of genes in the experimental data set which were significantly increased. EXPECTED, number of genes expected to be represented in the given pathway based on number of genes in gene set and in annotation pathway. OVER/UNDER, + for overrepresentations and – for underrepresentations. FOLD ENRICHMENT, fold increase in genes in set over expected.

**Supplemental Table 3. GO pathway analysis for Biological Process, Cellular Component, Molecular Function, and Reactome Pathways for pathways which are significantly upregulated in *Mcu*^CRISPR-KO^ 2838 clones over *Mcu*^WT^.** All genes significantly upregulated in *Mcu*^WT^ (p. adj.<0.05) were used for GO analysis. REFLIST, total number of genes in given annotation data set. GENESET, total number of genes in the experimental data set which were significantly increased. EXPECTED, number of genes expected to be represented in the given pathway based on number of genes in gene set and in annotation pathway. OVER/UNDER, + for overrepresentations and – for underrepresentations. FOLD ENRICHMENT, fold increase in genes in set over expected.

**Supplemental Table 4. GSEA results for Biological Process, Cellular Component, Molecular Function, and Hallmarks Pathways for pathways which are significantly upregulated in *Mcu*^WT^ 2838 clones over *Mcu*^CRISPR-KO^.** All gene sets with FDR < 0.25 or the top 10 gene sets, whichever is greater, are shown. SET, gene annotation set queried. NAME, pathway/data set name. SIZE, number of genes in set. ES, enrichment score. NES, normalized enrichment score. NOM p-value, nominal p-value. FDR q-value, false discovery rate q-value. RANK AT MAX, position in rank-ordered gene list at which maximum enrichment occurs. LEADING EDGE statistics: Tags – percent gene list before enrichment (surrogate readout for % of genes driving enrichment). List – percent of genes in the ranked list before the maximum enrichment score (indication of where maximum enrichment peak is located). Signal – Enrichment signal strength.

**Supplemental Table 5. GSEA results for Biological Process, Cellular Component, Molecular Function, and Hallmarks Pathways for pathways which are significantly upregulated in *Mcu*^CRISPR-KO^ 2838 clones over *Mcu*^WT^.** All gene sets with FDR < 0.25 or the top 10 gene sets, whichever is greater, are shown. SET, gene annotation set queried. NAME, pathway/data set name. SIZE, number of genes in set. ES, enrichment score. NES, normalized enrichment score. NOM p-value, nominal p-value. FDR q-value, false discovery rate q-value. RANK AT MAX, position in rank-ordered gene list at which maximum enrichment occurs. LEADING EDGE statistics: Tags – percent gene list before enrichment (surrogate readout for % of genes driving enrichment). List – percent of genes in the ranked list before the maximum enrichment score (indication of where maximum enrichment peak is located). Signal – Enrichment signal strength.

